# Programming animal physiology and behaviors through engineered bacteria

**DOI:** 10.1101/2020.08.15.232637

**Authors:** Baizhen Gao, Qing Sun

## Abstract

A central goal of synthetic biology is to predictably and efficiently reprogram living systems to perform computations and carry out specific biological tasks. Although there have been many advances in the bio-computational design of living systems, these advances have mainly been applied to microorganisms or cell lines; programming animal behavior and altering animal physiology remain challenges for synthetic biology because of the system complexity. Here, we present a bacteria-animal symbiont system in which engineered bacteria recognize external signals and modulate animal gene expression, twitching behavior, and fat metabolism through RNA interference. By using genetic circuits in bacteria to control RNA expression, we programmed the physiology and behavior of the model animal *Caenorhabditis elegans* with logic gates. We anticipate that engineered bacteria can be used more extensively to program animal metabolism and behaviors for agricultural, therapeutic, and basic science applications.

**One Sentence Summary:** Animal physiology and behaviors are programmed via engineered bacteria.

## Introduction

Recent advances in synthetic biology have resulted in highly programmable bacteria and mammalian cells that perform specific functions^1^. At the bacterial level, researchers have created logic functions and memory in living bacteria for bacteria and community oscillation, programmed bacteria to respond to signals, automated the circuit design process, and expanded the repertoire of logic gates so that they can recognize a variety of signals, including light and nutrients^1-3^. At the level of mammalian cells, researchers have engineered CAR-T cells and other mammalian cell types with switching logic circuits to program cell performance^4-8^. Despite rapid advances and promising results in bio-computational design, the programming of higher organisms remains largely unexplored because the engineering of an entire organism requires highly complex and sometimes impossible tasks, such as the delivery of gene editing cascades^9-11^ or the engineering of whole embryos^12,13^. Nevertheless, such programming is worthwhile because it will yield a greater understanding than we have so far of biological phenomena emerge from genetic and cellular events. The ability to change animal including insects and worm physiology and behaviors would open up possibilities for agriculture, biomedical, and bioenvironmental applications.

Animals share a life-long relationship with microbes, which play significant roles in animal nutrition, immunity, behavior, and metabolism^14-17^. Here, using bacteria as an engineering platform, we present an efficient strategy to program *Caenorhabditis elegans* GFP expression, twitching behaviors, and fat metabolism by building synthetic logic circuits in bacteria. We used the nematode *C*. *elegans* as the animal host as it has been extensively used as a model system to elucidate mechanisms of interaction between prokaryotes and their hosts^18,19^. The information from bacteria was transferred to *C*. *elegans* through an orthogonal gene transfer called RNA interference (RNAi), a widely used technique in which double-stranded RNA is exogenously introduced into an organism, causing knockdown of a target gene^20^. In *C*. *elegans*, RNAi is particularly easy to implement because it can be delivered by feeding the worms bacteria that express double-stranded RNA complementary to a *C*. *elegans* gene of interest^21^. Using this platform as well as metabolite-based host-bacteria interactions, the programming of logic gates on the organism levels could enable exquisite control of biological events in insects, mammals, or humans, for any number of applications.

We have engineered bacteria to produce RNA, and by putting the RNA expression under the control of genetic circuits in bacteria, we programmed *C*. *elegans* green fluorescence protein (GFP) expression profiles with “AND” and “OR” logic gates. We also achieved the programming of twitching behaviors and worm fat storage by placing the RNA targets under the control of logic gates. Overall, our results indicate that by taking advantage of bacteria-animal symbiosis, we can program animal physiology and behaviors that respond to environmental signals. This study thus presents a platform for the programming of higher organisms by harnessing engineered prokaryotic species to exert effects on eukaryotic organisms (Fig. 1).

**Figure 1.**
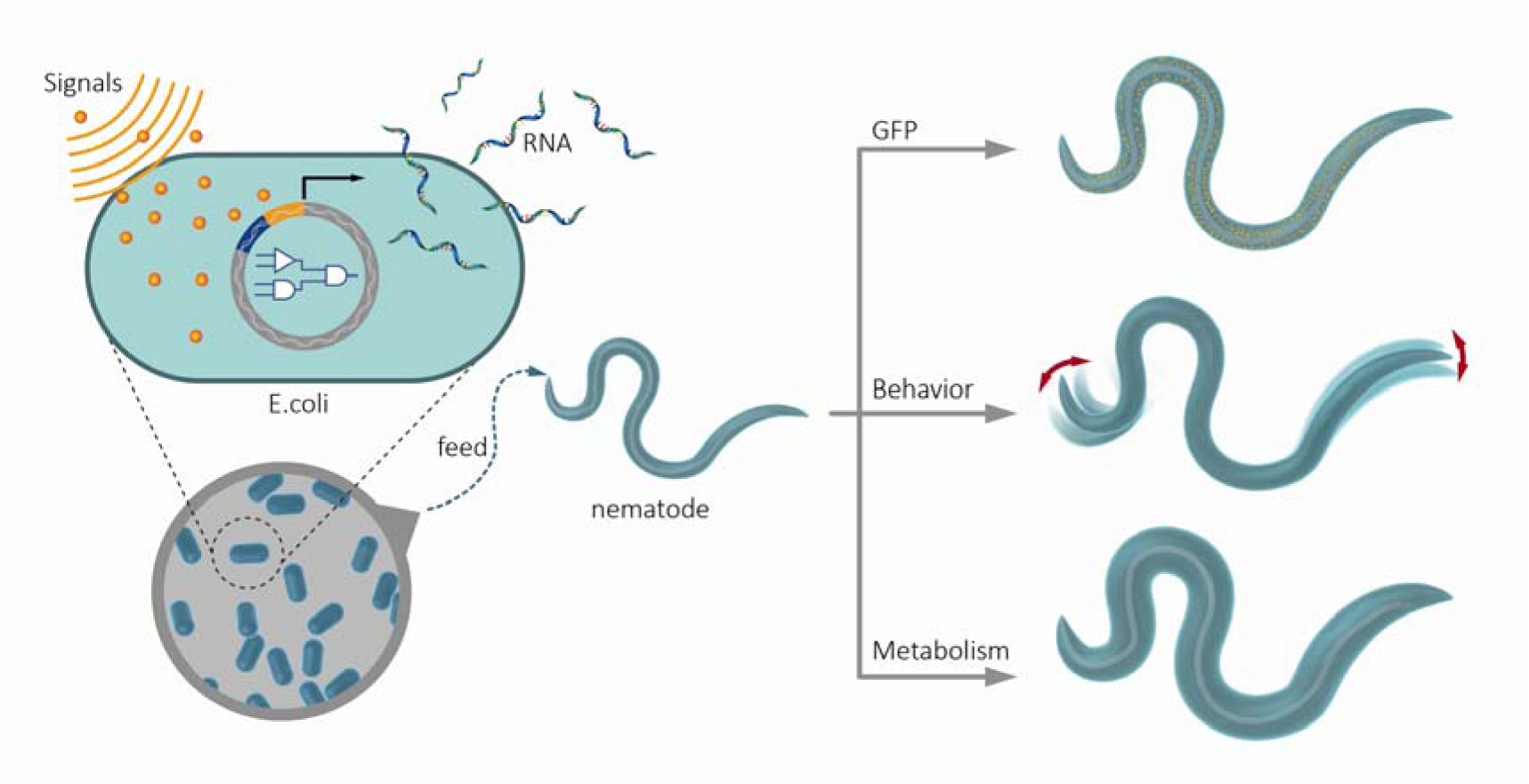
Engineering bacteria to program *C*. *elegans* GFP expression, behavior, and metabolism.

## Results

### Using RNA in ingested bacteria to silence GFP in *C*. *elegans* through RNAi

We started with the genetically engineered strain *C*. *elegans* SD1084, which expresses nuclear-localized SUR-5::GFP^22^. We intended to use the chemical inducer IPTG to silence GFP expression in this *C*. *elegans* strain through the worms’ ingestion of engineered bacteria. To achieve that, we put GFP transcription under the control of a bidirectional lac promoter, which induces the synthesis of RNA in *E*. *coli* under IPTG induction. By feeding these induced bacteria to *C*. *elegans*, we aimed to trigger GFP RNAi, which silences GFP in *C*. *elegans*^22^ (Fig. 2a). We expected that the differences between the “ON” and “OFF” states of the RNAi in *C*. *elegans* would be large enough for us to measure. However, the first round of experiments with full-length dsGFP led to GFP silencing under both the induced (i.e., with IPTG) and the uninduced (i.e., without IPTG) conditions (Fig. S1). We hypothesized that the leaky lac promoter already produced enough dsRNA to silence the worm GFP without IPTG. Rather than solve the problem by tuning the strength of the lac promoter, which is a less desirable approach because tuning all promoters for genetic circuits in the future is highly laborious, we decided to work on the intermediates between the bacteria and the worm to achieve an appropriate dynamic range.

**Figure 2.**
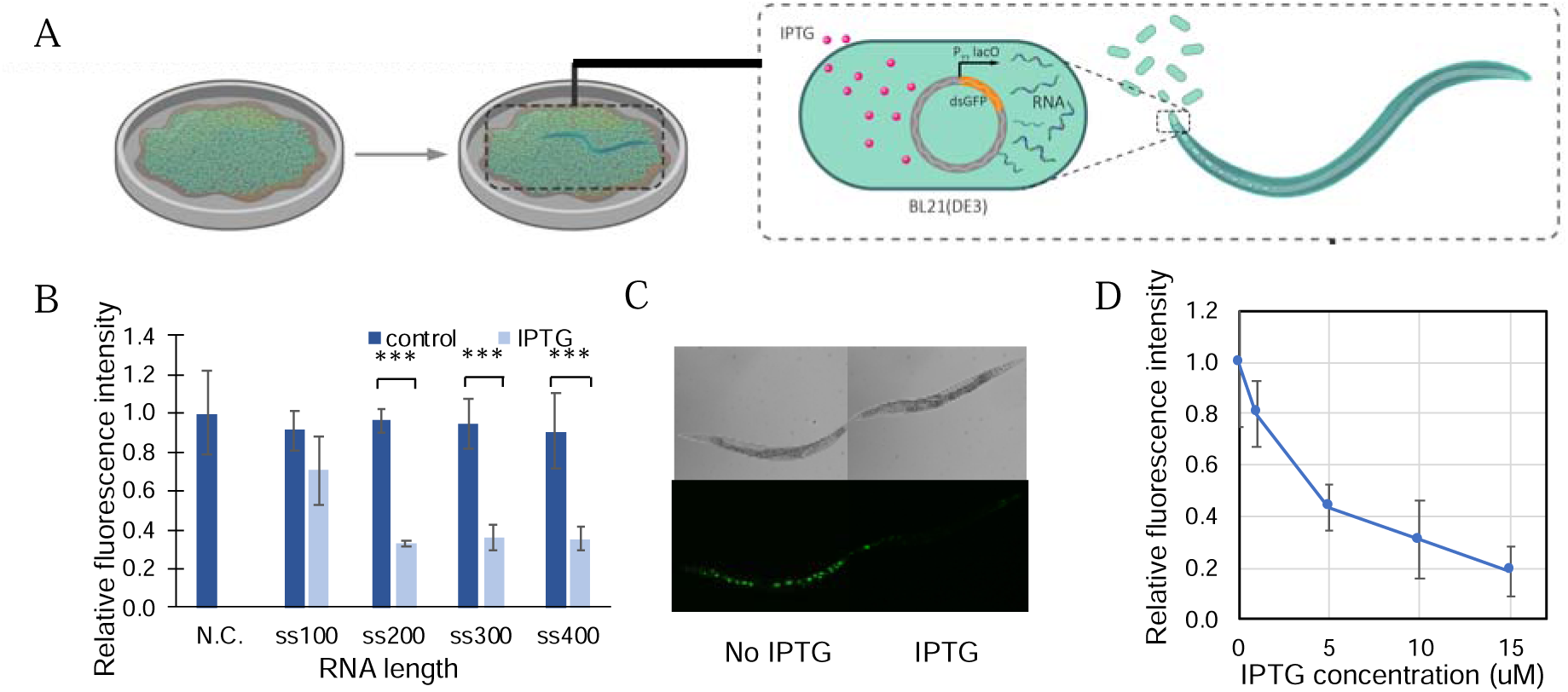
Using RNA in ingested bacteria to silence GFP in *C*. *elegans*. (A) Schematic of engineered *E*. *coli* triggering RNAi in *C*. *elegans* through feeding. (B) Quantification of *C*. *elegans* GFP fluorescence intensity controlled by bacteria yielded single-stranded GFP RNA of various lengths. (C) Optical and fluorescent images of *C*. *elegans* SD1084 GFP expression programmed by bacterial single-stranded 200 bp GFP RNA. (D) *C*. *elegans* SD1084 GFP intensity change caused by varying IPTG concentration-induced RNA synthesis. (***P < 0.001 Student’s t-test)

Systemic RNAi in *C*. *elegans* requires systemic RNA interference deficiency-1 (SID-1) protein for the transport of RNA^23,24^. SID-1 has been shown to have a higher binding affinity for longer RNA sequences^23^. We hypothesized that we could achieve the right dynamic range of RNAi by reducing the length of the RNA. We constructed plasmids for RNA synthesis with lengths ranging from 100 bp to 400 bp in double-stranded as well as single-stranded format (Fig. S2). After feeding the bacteria containing these plasmids to the worms, we found that single-stranded RNA sequences of 200bp, 300bp, and 400 bp achieved distinguishable “on” and “off” status of GFP expression in *C*. *elegans* with and without IPTG (Fig. 2B). We picked the single-stranded 200bp GFP RNA for subsequent experiments (Fig. 2C).

The efficiency of RNAi can be manipulated by changing the RNA concentration^20,23^, which can be controlled by the concentration of inducers. Using the 200bp design, we tested the dynamic range of the RNAi. We achieved a silencing of the GFP in *C*. *elegans* along a gradient, such that 81% of GFP in *C*. *elegans* was silenced when 15 μM of IPTG was supplied on the plate (Fig. 2D, S3).

### Programming GFP expression with logic gates

Logic gates, the building blocks of genetic circuits, can process complex signals and trigger downstream reactions^25^. Here we put GFP expression under the control of an “AND” or an “OR” gate and transferred the programming languages to *C*. *elegans* through RNAi.

An “AND” gate can be built by utilizing the split T7 polymerase (T7 RNAP)^26^. T7 RNAP is a single-subunit RNA polymerase that drives transcription by acting on its cognate promoter, the T7 promoter PT7. By splitting T7 RNAP between amino acids 179 and 180, a transcriptional AND gate is created whereby both fragments of the split protein are needed to drive transcription from PT7^26^. We used two orthogonal promoters, pBAD and pTet, to control T7 RNAP N-terminal and C-terminal expression, respectively. Both L-arabinose (Ara) and anhydrotetracycline (aTc), the respective inducers of these promoters, are needed to trigger GFP transcription in the bacteria and, in turn, RNAi expression in the worms. With this system, worm GFP expression was silenced by more than 70% in the presence of both signals (Ara and aTc) whereas GFP was still expressed (more than 95%) with either no signal or only one of the two signals (Fig. 3A, S4A).

**Figure 3.**
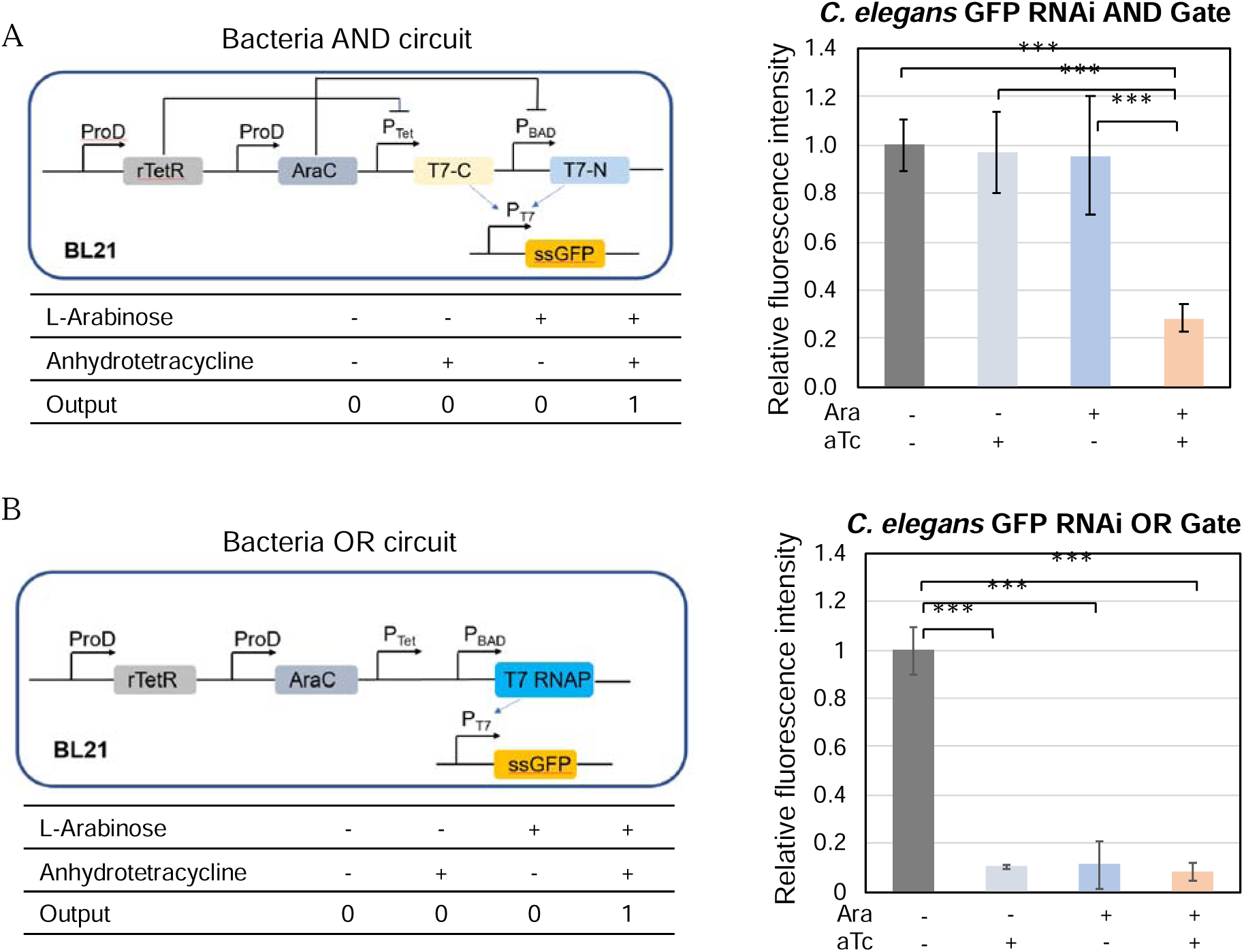
GFP expression in *C*. *elegans* programmed by AND and OR gates. Single-stranded 200bp GFP RNA synthesis in *E*. *coli* BL21, controlled by (A) AND gate or (B) OR gate, interferes with GFP expression in *C*. *elegans* that have ingested the bacteria. (***P < 0.001 Student’s t-test)

For the OR gate, the pTet and pBAD were arranged in tandem to control the synthesis of full-length T7 polymerase, which transcribes GFP under the control of promoter PT7. Activation of either pTet or pBAD was expected to lead to GFP silencing in *C*. *elegans*. We observed that the addition of one or both of the inducers led to over 90% reduction in *C*. *elegans* GFP intensity, while the no-inducer control group only had a 24% reduction (Fig. 3B, S4B).

### Programmed *C*. *elegans* behavior and metabolism with logic gates

After demonstrating that *C*. *elegans* GFP expression could be programmed with bacterial genetic circuits, we programmed essential gene silencing that affects *C*. *elegans* behaviors and metabolism. First, we programmed the twitching behavior in *C*. *elegans* by interfering with the expression of the unc-22 protein, which encodes a polypeptide involved in locomotion^24^. Using similar approaches as those described above for GFP, we screened out double-stranded 400bp and 200bp unc-22 for the “OFF” and “ON” status of worm twitching behavior (Fig. 4A). The twitching behavior was characterized by observing the *C*. *elegans* body movement under a microscope immediately after the worms were anesthetized by levamisole^27^. The AND gate was constructed by using the same genetic circuit-carrying plasmid and replacing the gene of interest with unc-22. Only the worms grown on plates with both inducers showed the twitching behavior. 100% of the worms exhibited twitching behavior in the presence of both signals while 0% of the worms on the other three plates (no inducers plate, aTc only plate, and Ara only plate) showed twitching behavior. Using the same OR gate but replacing the GFP in the OR genetic circuits with unc-22, 100% of the worms grown on plates with one or both inducers showed twitching behaviors (Figure 4B, SV1).

**Figure 4.**
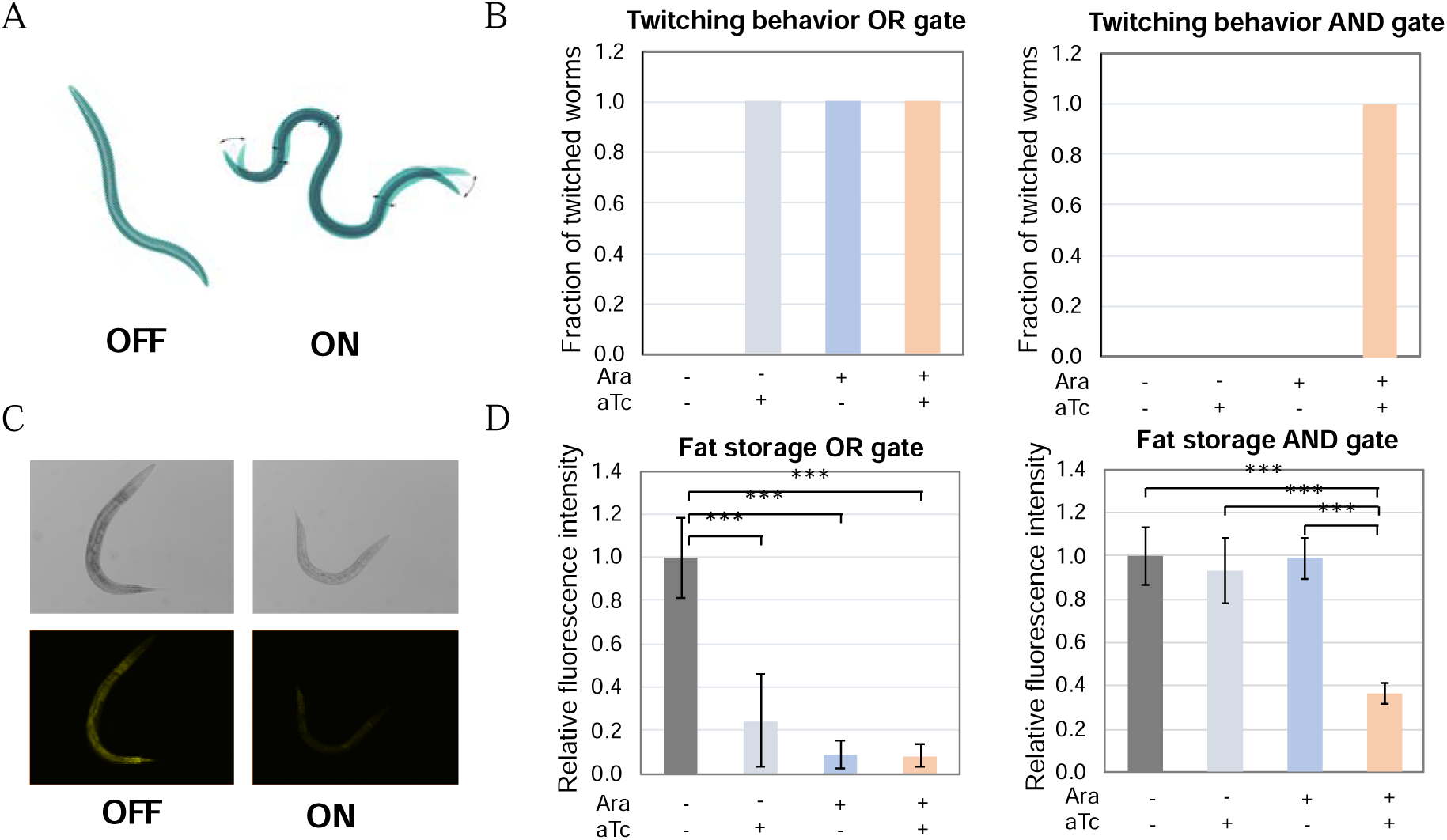
Worm twitching behavior and fat storage programming through engineered bacteria. (A) Schematic of twitching behavior RNAi in *C*. *elegans*. (B) *C*. *elegans* twitching behavior in response to AND and OR gate programming through bacteria expressing unc-22 RNA (C) Reduced fat storage due to sbp-1 silencing, indicated by Nile Red staining (D) AND and OR gate-controlled *C*. *elegans* metabolism through sbp-1 RNA. (***P < 0.001 Student’s t-test)

Besides twitching behavior, we programmed *C*. *elegans* metabolism through the sbp-1 gene. Previous studies have shown that sbp-1 facilitates fat storage, and silencing sbp-1 using RNAi leads to reduced body fat storage in *C*. *elegans*^28^. We modulated *C*. *elegans* fat storage by using sbp-1 RNAi (Fig. 4C); the modulation of fat storage was characterized by changes in Nile red staining. When the worms grew on plates with bacteria containing plasmids for the AND gate and sbp-1 RNA synthesis, only the group that had both inducers showed a significant reduction in both worm body size and fat storage as indicated by a reduction in Nile red staining. For the engineered OR gate, the group without inducers had a body size and a level of fat storage similar to those of normal *C*. *elegans* (Figure 4D, S5). All other groups, which were exposed to either one or both of the inducers, showed an over 60% reduction in fat storage, demonstrating the successful OR gate programming of *C*. *elegans* fat storage.

## Discussion

Although the technology for genetically engineering living organisms has advanced rapidly, the complex design of genetic circuits and gene editing in animals remain challenging because of the complexity of these systems. In contrast, genetic circuit design and editing have been highly successful in bacteria and mammalian cells. Bacteria affect the metabolism, immunity, and behavior of the animals they live in and on. We took advantage of the symbiosis between bacteria and animals to engineer prokaryotic cells as an indirect, alternative approach to programming animal physiology and behaviors.

Using *E*. *coli* as the engineering platform and *C*. *elegans* as the host animal, we tuned *C*. *elegans* gene expression by changing the quantity of the RNA produced by the bacteria. The silencing of *C*. *elegans* gene expression was accomplished, through RNAi, with a reasonable dynamic range of programmable gene expression, behavior, and metabolism. We determined the effective dynamic range by screening out different lengths of RNA as mediators between bacterial and animal. An alternative way to tune the dynamic range is by changing the inducer concentration and incorporating T7 lysozyme to reduce basal RNA expression. This approach for basal level reduction as an alternative to shortening the RNA length is important because the logic gates can be tuned for enzyme expression rather than RNA synthesis. Furthermore, we applied genetic circuits to program *C*. *elegans* GFP expression, twitching behaviors, and fat storage through “AND” and “OR” logic gates. The fact that none of these worm phenotypes were affected by the inducers used in this study (L-arabinose and anhydrotetracycline) demonstrated that *C*. *elegans* underwent real programming by the engineered bacteria. The most striking advantage of this system is the ease with which system complexity can be extended by applying more complicated logic gates. Since bacteria have already been engineered with more complicated logic gates, memory, and sensing more signals, we should be able to program the animals with three or four input logic gates^3^ or extend the capability of animals to recognize external signals^2,29,30^ transmitted by these engineered bacteria.

RNAi is especially useful in the area of pests control and worm behavior interference as a gene silencing tool. However, for higher organisms in which RNAi would not be sufficient to alter characteristics or behavior, bacterial metabolites could be exploited to serve as mediators between bacterial and animal (e.g., bile acid^31-33^, butyrate^34-36^, colonic acid^37^, nitrite^38-40^) to communicate between bacteria and higher organisms. Metabolites as intermediaries have broad applications for studying microbe-host symbiosis *in situ*; for example, they can be used as a drug delivery system. Indeed, bacterial metabolites operate as signaling molecules in the microbiome. Secondly, in this study we used synthetic inducers that were manually introduced into the host environment. By engineering bacteria to respond to the host’s native signals, usually in the form of nutrients (sugar^29,30^ and hormones^41,42^), similar programming can be applied for *in situ* detection and responses.

In summary, we have programmed *C*. *elegans* through the ingestion of engineered bacteria to exhibit changes in physiology, behavior, and metabolism. Our strategy could be extended to other bacteria or hosts, including fruit flies, zebrafish, plants, or mammals. The potential to program higher organisms using bacterial RNA or metabolites as intermediaries enabled us to study the bacteria-animal interaction. Our system can be developed for applications to pest control, disease diagnosis, and the delivery of therapeutics.

## Acknowledgments

QS conceived of the study. BG and QS designed experiments. BG constructed plasmids and strains and conducted all the worm-related experiments. QS and BG wrote the manuscript. The authors declare no competing interests.

## Materials and Methods

### Bacterial strains, worm strains, and growth conditions

Bacterial strain *E*. *coli* BL21 (DE3) was used for testing of constructed plasmids encoding the RNA production without logic gate control. *E*. *coli* BL21 was transformed with both the constructed logic gate and RNA production plasmids. *E*. *coli* OP50 was used for maintenance of *C*. *elegans*. Bacterial cultures were grown in LB media with appropriate antibiotics in a 37°C shaking incubator overnight before use.

Wild-type *Caenorhabditis elegans* Bristol N2 and SD1084 (Caenorhabditis Genetics Center, Minnesota, USA) were cultured at room temperature on NGM (Nematode Growth Medium) agar plated with *E*. *coli* OP50 as described in WormBook^43^. Synchronized L1 worms were used in all RNAi treatment experiments. Synchronization was performed following the standard protocol^43^. Briefly, worms were allowed to grow on NGM plates with *E*. *coli* OP50 for 2-4 days then washed off the plates to collect a large number of gravid hermaphrodites, and eggs were isolated by bleach-NaOH lysis of the gravid worms. After 4X wash in M9 buffer to remove the residual bleach and NaOH, eggs were allowed to hatch in 10 mL of M9 buffer overnight and develop into starved L1, which were then used for the RNAi experiments.

### Plasmid construction for RNA synthesis and logic gates

All plasmids were constructed using PCR and Gibson assembly. Gene sequences used in this study are detailed in Table S1. For double-stranded GFP and unc-22 RNA synthesis, the RNA sequence of interest was inserted between two T7/LacO for the synthesis of both complementary strands on a pET24a vector. (Using this method, pET24a-T7-dsGFP, pET24a-T7-dsGFP-100bp, ET24a-T7-dsGFP-400bp, pET24a-T7-dsunc-22-200bp, and pET24a-T7-dsunc-22-400bp were constructed.) Additional T7 lysozyme and LacI were constitutively expressed under the control of the proD promoter for double-stranded sbp-1 RNAi to reduce background expression level for the construction of the plasmid pET24a-dssbp-1-proD-lysS-proD-lacI.

Plasmids for single-stranded RNA synthesis were constructed by putting the RNA sequence of interest downstream of one T7/Lac on a pET24a vector. (Plasmids pET24a-T7-ssGFP-100bp and pET24a-T7-ssGFP-400bp were synthesized this way)

The AND gate plasmid (pTet-Ara-Split-T7-AND-Gate-proD) was modified from plasmid pTSlb-wt by constitutively expressing the TetR and araC repressors under proD promoters. The C-terminus of T7 RNA polymerase was regulated by pBAD, and the N-terminus of the T7 RNA polymerase was regulated by pTet. The OR gate (pTet-Ara-OR-Gate-proD) was constructed similarly by putting the full-length T7 RNA polymerase under the control of the two promoters arranged (pBAD and pTet) in tandem.

### Feeding *C*. *elegans* with the engineered bacteria for RNAi

*E*. *coli* BL21 bacteria transformed with plasmids for RNA synthesis and logic gates were grown at 37°C in LB with 50 μg/ml kanamycin and 25 μg/ml chloramphenicol, then seeded onto NGM plates supplemented with appropriate inducers as indicated in the supplementary information. After overnight growth of the seeded bacteria on NGM plates, synchronized L1 larvae were added to the plates and day-1 adult worms were used for phenotype assays after 2 days.

To program *C*. *elegans* GFP expression using IPTG, plasmids pET24a-T7-dsGFP, pET24a-T7-dsGFP-100bp to pET24a-T7-dsGFP-400bp were transformed into *E*. *coli* BL21(DE3) separately and fed to the worms. To induce the RNA synthesis in *E*. *coli*, 1 mM of IPTG was added into the NGM plates. To program GFP expression with an AND gate, *E*. *coli* BL21 was co-transformed with pTet-Ara-Split-T7-AND-Gate-proD and pET24a-T7-ssGFP-200bp, and pTet-Ara-OR-Gate-proD and pET24a-T7-ssGFP-200bp were co-transformed for OR gate programming. L-arabinose and anhydrotetracycline at 0.2 mg/mL and 0.1 μg/mL, respectively, were used as the external signal for both AND and OR gates.

For twitching behavior AND gate programming, *E*. *coli* BL21 was co-transformed with pTet-Ara-Split-T7-AND-Gate-proD and pET24a-dsunc-22-400bp, and RNA synthesis was induced by 2 μg/mL L-arabinose and 0.1 μg/mL anhydrotetracycline. To program twitching behavior using an OR gate, *E*. *coli* BL21 was co-transformed with pTet-Ara-OR-Gate-proD and pET24a-dsunc-22-200bp, and expression was induced by 0.2 mg/mL L-arabinose and 0.1 μg/mL anhydrotetracycline.

AND gate programming of fat storage in *C*. *elegans* was achieved by co-transforming *E*. *coli* BL21 with pTet-Ara-Split-T7-AND-Gate-proD and pET24a-dssbp-1-proD-lysS-proD-lacI, and expression was induced by 2 μg/mL L-arabinose and 0.1 μg/mL anhydrotetracycline. OR gate programming of fat storage was done by feeding the worms with *E*. *coli* BL21 co-transformed with pTet-Ara-OR-Gate-proD and pET24a-dssbp-1-proD-lysS-proD-lacI, and induced by 0.2 mg/mL L-arabinose and 0.1 μg/mL anhydrotetracycline.

### *C*. *elegans* GFP fluorescence imaging

Day-1 adult worms were rinsed off the NGM plates using PBS+0.01% Triton X-100 (PBST) and washed twice with PBST to remove the bacteria. The worms were then transferred onto glass slides with 2% agarose containing 1mM levamisole. Fluorescence in *C*. *elegans* was imaged on an Axiovert 200M fluorescence microscope using the FTIC filter.

### Nile red staining

*C*. *elegans* fat storage was characterized by Nile Red staining as previously described with slight changes^44^. Synchronized L1 stage N2 worms were put onto the bacteria lawns, and allowed to grow for 2 days at room temperature. Young adult worms were washed off the plates by PBST and then fixed by 40% isopropanol (v/v) for 3 minutes. For lipid staining, a Nile Red stock solution of 0.5 mg/ml in acetone was first prepared. The working solution was then made by adding 6 μL of the stock solution into 1mL of 40% isopropanol. After the worms were fixed and the supernatant removed, the working solution was added, and the worms were incubated in the working solution in the dark for 2 hours. Following the 2-hour incubation, the supernatant was again removed, 150 μL PBST was added, and the worms were incubated for another 30 minutes. The worms were then imaged using an Axiovert 200M fluorescence microscope, and fluorescence intensity was quantified in ImageJ.

### Twitching behavior recording

After *C*. *elegans* were fed with the control and engineered *E*. *coli* for 2 days, the worms were washed off the plates using PBST. The worms were then washed once with PBST and resuspended with 20 μL 3mM levamisole (in water)^27^. Resuspended worms were immediately transferred onto glass slides for recording under a microscope^27^.

